# *Anopheles gambiae* maternal age and parous state control offspring susceptibility to *Plasmodium falciparum*

**DOI:** 10.1101/2020.01.27.922070

**Authors:** Christian Mitri, Isabelle Thiery, Marie-Thérèse Lecoq, Catherine Thouvenot, Solange Touron, Annie Landier, Emmanuel Bischoff, Catherine Bourgouin

## Abstract

Maternal effects have been reported in many organisms whereby exposure to environmental stress, either toxics or pathogens will impact on progeny response to these stresses. Here we show that *Anopheles gambiae* susceptibility to *Plasmodium falciparum* is dependent upon maternal effects driven by females not previously exposed to the parasite. The maternal effect involved both mother age and reproductive state. Offspring of old females or from a 4^th^ gonotrophic cycle are more susceptible than offspring from young females. These maternal effects also contribute to overall better fitness of the offspring. As mosquito population age structure contributes heavily shaping malaria transmission, consequences of this novel finding should be taken into account in further strategies for controlling malaria transmission.

## Main Text

Malaria transmission is dependent on the bioecology and bionomics of *Anopheles* mosquitoes. Malaria transmission modelling relies on estimating mosquito vectorial capacity C (*1, 2*) which includes estimation of several entomological indices such as the number of mosquitoes, their daily biting rate and the extrinsic incubation period that quantifies the length of parasite development within the mosquito before it can be transmitted. One parameter that is not clearly defined in the estimation of C is vector competence, which corresponds to the ability of a mosquito to sustain *Plasmodium* development up to the production of infective sporozoites in her salivary glands (*3, 4*). Because difficult to measure, this parameter has often been overlooked and the competence of *An. gambiae* to *P. falciparum* has long been assumed to be identical among populations in malaria transmission modelling. Advances in genomics and genetics showed that wild *An. gambiae* populations are indeed composed of genotypes controlling refractoriness or susceptibility to *P. falciparum (5, 6)*. The penetrance of those phenotypes varies and quantitative genetic approaches identified several QTL contributing to these refractory/susceptible phenotypes. Within these QTL much attention has been given to innate immunity genes (*5, 7*). On the other hand, there is accumulating evidence that vector competence is also dependent upon genotype by genotype interactions involving both vector and pathogen genotypes (*8–10*).

Vectorial capacity reflects several life history traits of a vector population, such as her reproduction, or parous, state, age and survival rate. Each of these traits contribute to shaping population structure which is one the most sensitive parameters influencing the epidemiology of vector-borne diseases (*3, 11*). Reducing any of these parameters will obviously reduce pathogen transmission and this has been exemplified in the last decade in *Aedes* mosquitoes and Dengue virus transmission through shortening vector survival (*12–16*). Impact of mosquito life history traits on vector competence has recently gained interest, focusing mostly on larval development, mosquito size, fitness and receptivity to malaria parasite or arbovirus (*4, 17–24*). One study showed that *An. gambiae* exposed during their larval life to pathogens will produce females with reduce susceptibility to a rodent *Plasmodium (25).* In various systems, maternal exposure to toxics or stresses such as food quality or deprivation can drive phenotypic changes in the offspring that are collectively referred as maternal effects (*26, 27*). Along this line, exposing *Anopheles* female mosquitoes to *P. falciparum* produced offspring with reduced susceptibility to this parasite species (*25, 28*). In Drosophila and in Humans, several studies addressed the impact of mother age on offspring fitness and/or susceptibility to diseases (Bell1918 in (*29*), (*29, 30*). We investigated here the concept of maternal effect on competence of *An. gambiae* to *P. falciparum* by looking at two essential parameters shaping malaria transmission: the age of the mothers and their gonotrophic status (parity) on susceptibility of the offspring to *P. falciparum*.

We first analysed the receptivity to *P. falciparum* of four successive offspring born from the same mothers. These offspring or daughter populations are thereafter named DP1 to DP4 (Figure 1A). Mosquitoes were raised in such a way that adults from successive egg collections of the same mothers, reared separately, will emerge at the same time. At the peak of emergence, pools of 100 females from each DP were selected on the same day, and fed on the same batch of *P. falciparum* gametocytes, five days later. Receptivity of each DP to *P. falciparum* was measured as the proportion of infected females (infection prevalence) on day 7 post infection and as the number of developing parasites in each infected female (infection intensity). Pooled results from two independent replicates clearly show that the proportion of infected females increased gradually and significantly from DP1 to DP4, with DP4 being fully susceptible (Figure 2A, SupTable1). Furthermore, the number of oocysts developing in DP4 was significantly higher than that developing in the 3 other DPs (Figure 2B, Sup Table2). All DPs have the same mothers but differ by mother age at birth (see Fig1A). Therefore, our results suggest that either females born from old mothers are more susceptible to *P. falciparum* than females born from younger females, and/or the physiological parous state of the mothers, having produced more than one offspring, imprints on adult offspring receptivity to *Plasmodium*. Producing the four DPs to be infected on the same gametocyte populations, imposed that larval development duration differed over 10 days between DP1 and DP4 (Fig 1A). As age at pupation impacts on adult mosquito immune-competence (*31*) the observed difference in susceptibility among the four DPs might also be a consequence of fitness differences acquired during the larval stages. To test this hypothesis, DP1 and DP4 were produced from different mother populations to accommodate for similar larval development duration. The experimental design was such that DP1 and DP4 were assessed for *Plasmodium* susceptibility on the same gametocyte populations (Fig 1B). Pooled results from 5 independent biological replicates show that in those conditions DP4 was again more susceptible than DP1, with high statistical significance for both infection prevalence and parasite loads (Figure 3). Therefore, larval development duration does not contribute as a major determinant to the difference in *Plasmodium* receptivity/susceptibility between DP1 and DP4.

**Figure 1:**
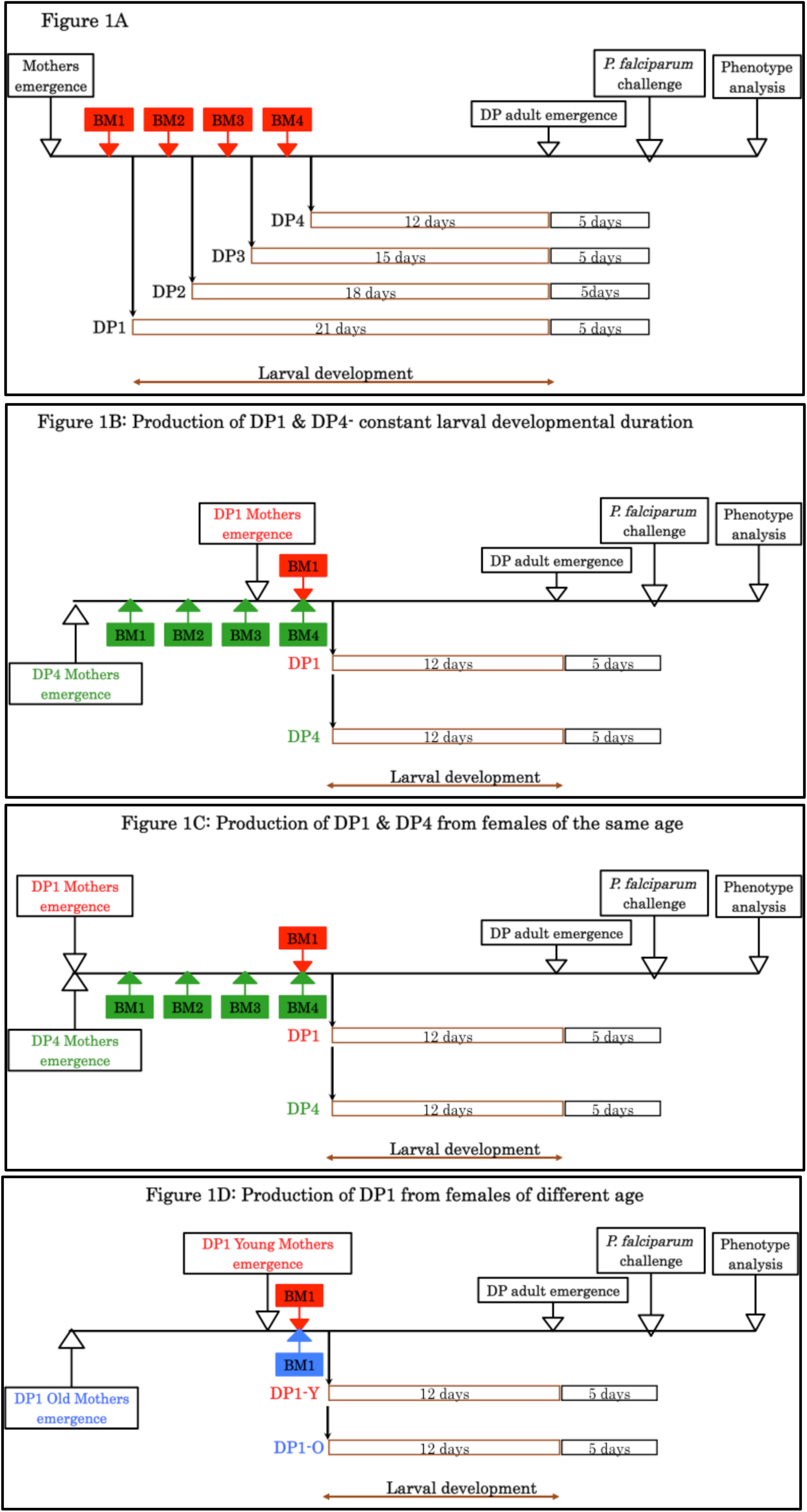
Schematic representation of *Anopheles gambiae* progeny production and infection with *Plasmodium falciparum* NF54. 1A): production of DP1, DP2, DP3 and DP4 from the same mothers. 1B): production of DP1 and DP4 from mothers of different age. 1C): production of DP1 and DP4 from mothers of the same age but having accomplished 1 or 4 gonotrophic cycles. 1D) Production of DP1 progenies from young mothers (=DP1-Y) and from old mothers (=DP1-O). DP = Daughter Progeny; BM= Blood meal.

**Figure 2:**
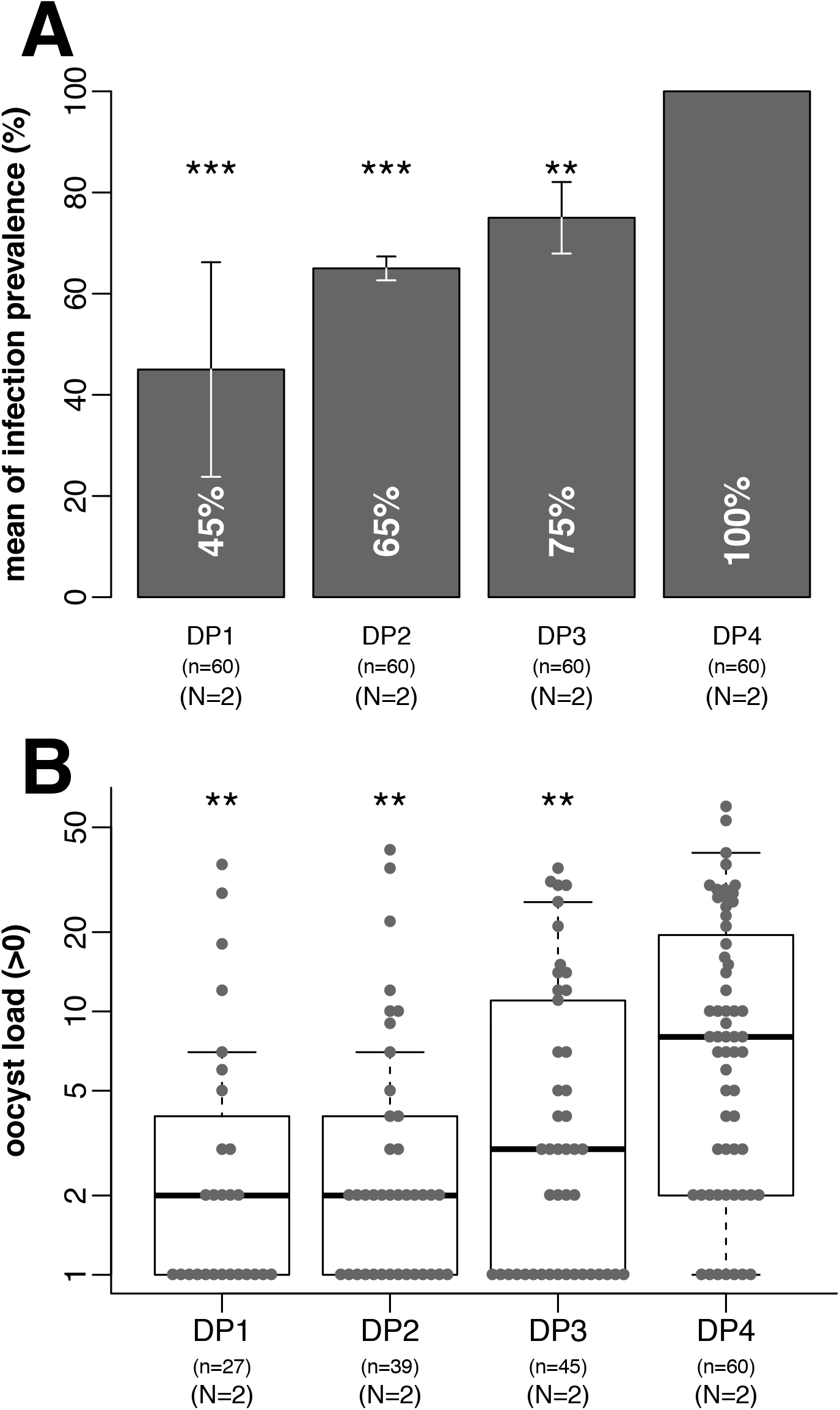
The number of gonotrophic cycles of the mother increases both prevalence (panel A) and intensity (panel B) of P. falciparum infection in the daughter populations (DP). Graph labels and statistical tests for this and subsequent figures: tests of infection prevalence indicate mean infection prevalence within histogram bars, error bars indicate standard error. For tests of infection intensity, the y-axis is logarithmic to clarify low-intensity effects, boxplots delineate the first and third quartile, median is indicated within the box, and error bars are 1.5 times the interquartile range. Sample sizes (N) indicate the number of independent replicate experiments, (n) the total number of mosquitoes dissected across replicates. Differences in infection prevalence were statistically tested using the Chi-Square test, and analysis of oocyst intensity used the Wilcoxon signed rank non-parametric test. Statistical differences in prevalence and intensity were first tested independently for each independent replicate as described above and p-values were empirically determined using 10^5 Monte-Carlo permutations. Following independent statistical tests, the p-values from independent tests of significance were combined using the meta-analytical approach of Fisher [1] when the direction of change of each independent replicate was concordant. Statistical tests have been done for all pairwise comparisons and p-values have been adjusted for multiple testing using the Bonferroni procedure. P-values reported on the plots correspond to the comparison of DP4 with the other daughter populations. P-values for all comparisons are reported in Supplemental Table 1 and 2 for prevalence and intensity, respectively. (Significance levels of adjusted Fisher-combined p-values: n.s., not significant; * p-value<0.05; ** p-value <0.01; *** p-value <0.001)

**Figure 3:**
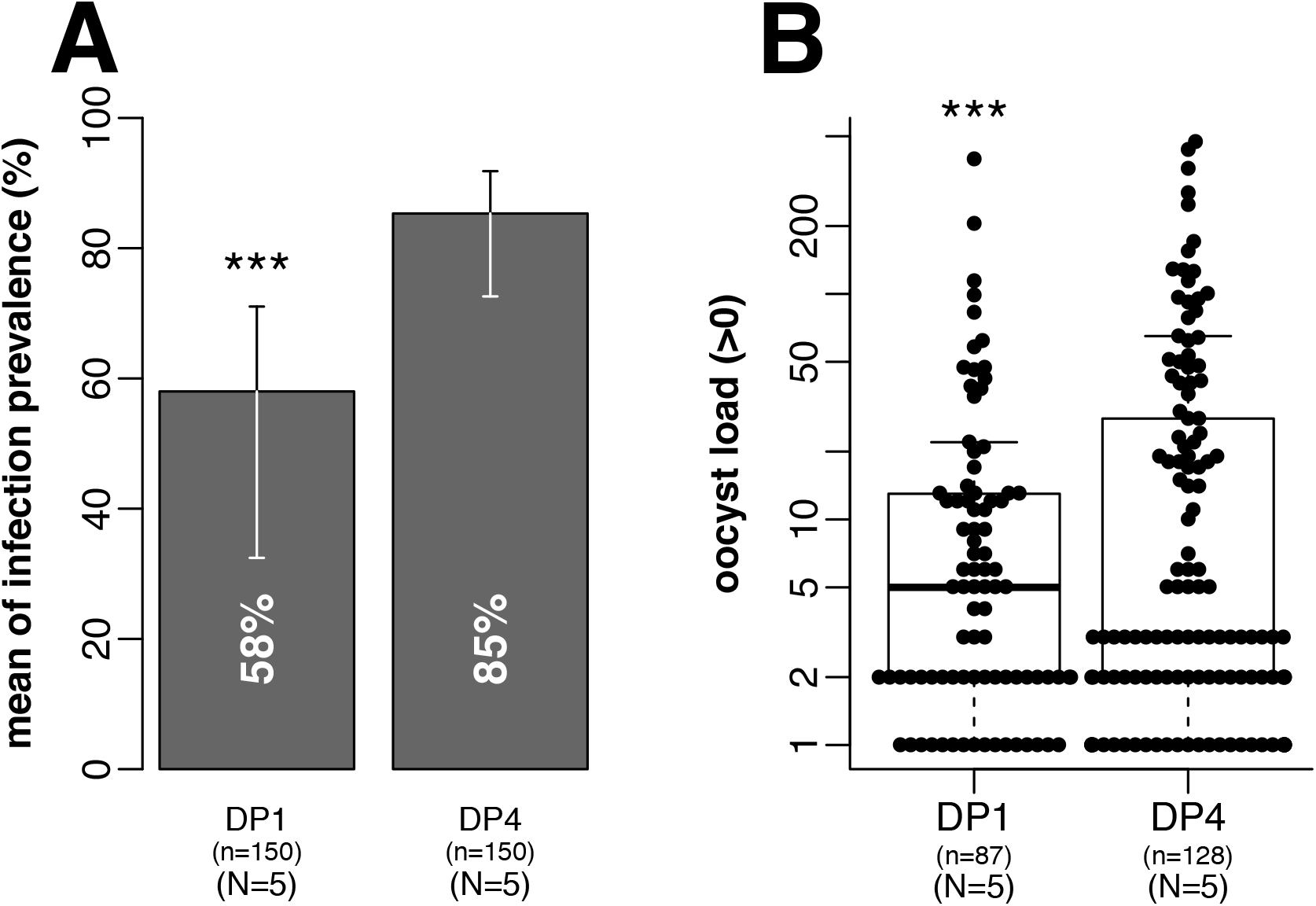
Combined mother age and number of gonotrophic cycles increases both prevalence (panel A) and intensity (panel B) of *P. falciparum* infection in the daughter populations (DP). DP1 and DP4 were produced as depicted in Figure 1B; (A) prevalence of infection. (B) intensity of infection. Graph labels and statistical tests as in Fig 2 legend. Note than n in panel B takes into account only sample harboring at least on oocyst. Significance levels of adjusted Fisher-combined p-values: n.s., not significant; * p-value<0.05; ** p-value <0.01; *** p-value <0.001).

Next, we determined whether the parous state of the mother, her age, or both, contribute to variation in offspring susceptibility to *Plasmodium*. First DP1 and DP4 were produced from females having the same age at egg laying (Fig 1C). Larval development duration was similar and the paired offspring were infected on the same *Plasmodium* gametocyte populations. Under these conditions, progeny born from a 4^th^ gonotrophic cycle (DP4) are more susceptible to *P. falciparum* than progeny born from a first gonotrophic cycle of females of the same age at egg laying (DP1) (Figure 4A). This is statistically significant for prevalence of infection only (Figure 4A and sup Fig1). Conversely, we produced DP1 from young (5 days old) and old females (15 days old) (Figure 1D) that were infected on the same *P. falciparum* gametocyte populations. DP1 from 15 days old females (DP1-O) were more susceptible to *P. falciparum* than DP1 from 5 days old females (DP1-Y) (Figure 4B and sup Fig1). This is valid for infection prevalence only and not for parasite loads, which were similar among the infected females (Figure 4B and sup Fig1).

**Figure 4.**
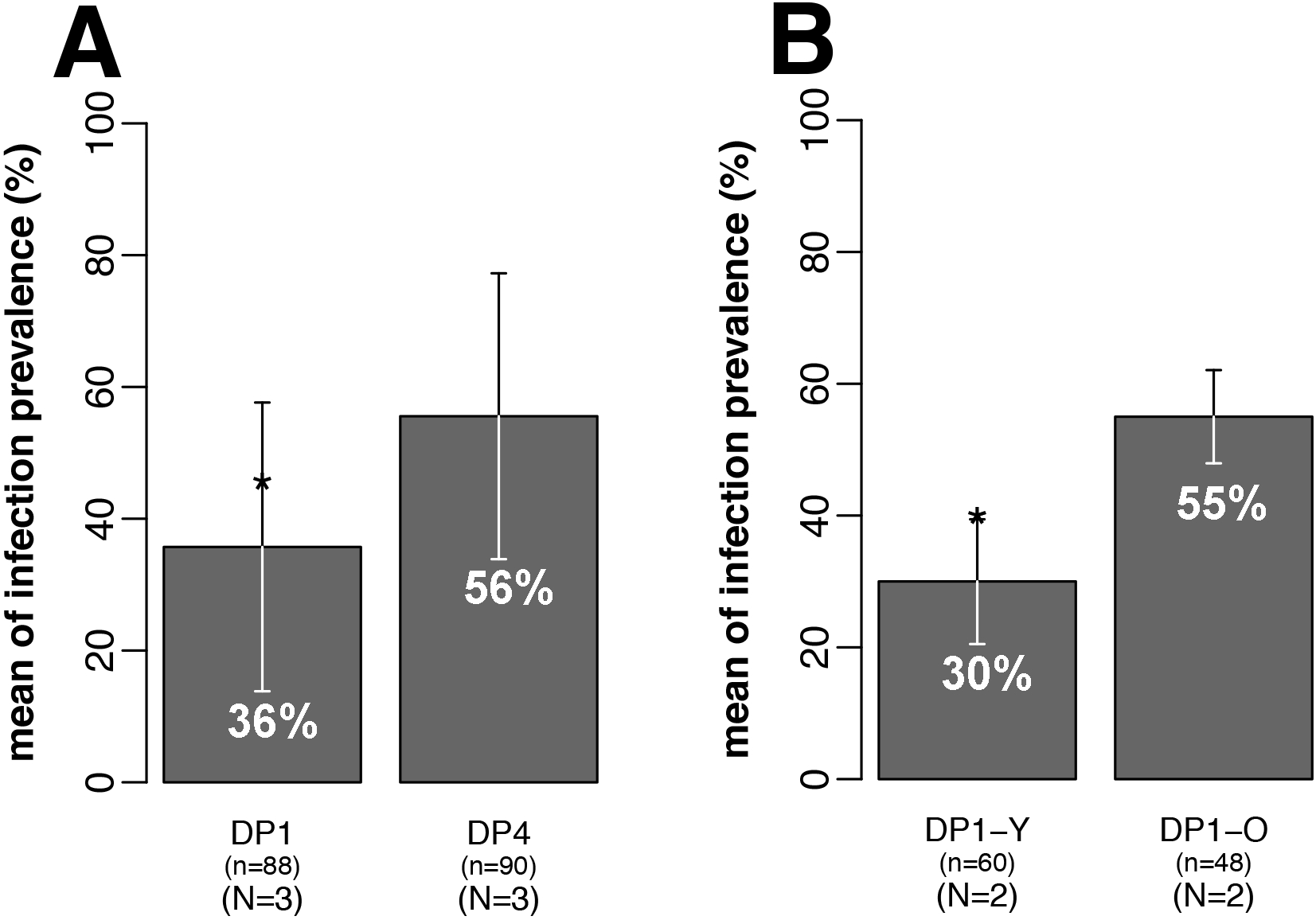
Similar contribution of gonotrophic cycles and mother age to increased receptivity to P. falciparum in the daughter populations (DP). Panel A: DP1 and DP4 were produced as depicted in Figure 1 C. Panel B: DP1 were produced form old or young females as depicted in Figure 1D. Graph labels and statistical tests as in Fig 2 legend. Significance levels of adjusted Fisher-combined p-values: n.s., not significant;* p-value<0.05; ** p-value <0.01; *** p-value <0.001)

Collectively, these results suggest that both the age and the parous state of the mosquito mothers contribute as major traits for offspring susceptibility to *P. falciparum*. Both these maternal history traits act on prevalence of infection and intensity of infection (oocyst load). However, when assessed individually they do not seem to contribute to differences in infection intensity. Comparison of those data with data presented in figure 2 suggests that larval development duration might also contribute to some extend to differences in progeny susceptibility to *P. falciparum*.

To determine whether maternal effects on progeny susceptibility to pathogens is restricted to *P. falciparum* or a global feature of *An. gambiae*, we next challenged DP1 and DP4 with *Beauveria bassiana*, an entomopathogenic eukaryote fungus (*32*). This fungus differs not only phylogenetically from the malaria parasite, but also in both its mode of infection (outside attack from the cuticle) and its pathogenic effect, as it kills the insect (*32*). DP1 and DP4 adults were produced as described in Fig 1B and exposed to *B. bassiana* spores (Method in sup data). Results from 2 independent experiments (Figure 5) show that exposed DP1 were killed faster than exposed DP4, and conversely that DP4 exhibits higher resistance to the fungus than DP1. This is in sharp contrast with higher susceptibility of DP4 to *P. falciparum*. We then assessed whether the differential survival characteristics observed between DP1 and DP4 infected with *B. bassiana*, might be due to fitness differences. Hence, we measured survival rates of DP1 and DP4, in absence of infection. Results from 2 biological replicates (Sup Figure 2) show that DP4 live significantly longer than DP1. Overall, these data indicate that offspring from old females (DP4) are more fitted in term of longevity than offspring of young females (DP1) independently of the infection status.

**Figure 5:**
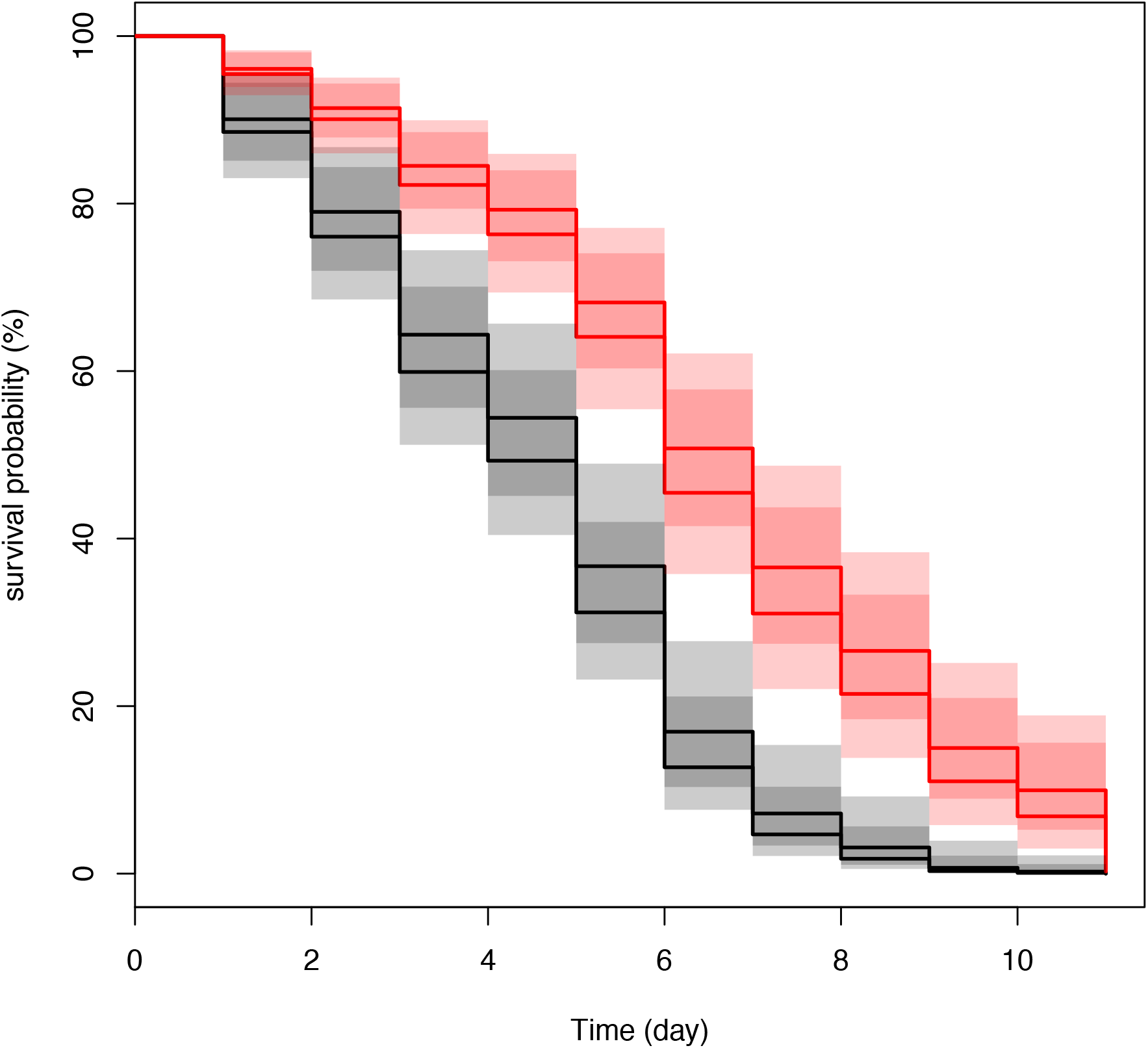
*An. gambiae* DP4 survived better than DP1 when exposed to *Beauveria bassiana*. Survival curves of *An. gambiae* DP1 (black) and DP4 (red) exposed to the spores of *B. bassiana*. DP1 and DP4 were produced as described in Figure 1B. Day 0 corresponds to the day of mosquito exposure to *B. bassiana*. Two replicates were performed using 50 female mosquitoes in each replicate and condition (DP1/DP4). A Cox proportional hazards regression model was fitted to the data using condition and replicate as predictor terms. P=2.7e^−09^.

Taken together, our findings constitute the first report of maternal effect in *An. gambiae* susceptibility to *P. falciparum* driven by mother age and parous state. Maternal effect on fitness of offspring has been reported in several systems from Daphnia to Human (*33, 34*). One of the major factor contributing to this phenomenon is environmental constraints, the mothers being exposition to toxics or limited access to food. The phenotypic pattern can be higher resistance to toxic of the offspring or shorten live in food deprivation context (Humans) or higher susceptibility to pathogens. In our work, none of these parameters can apply as all mother faced the same environmental and access to food conditions and were not exposed to *Plasmodium*. Age at reproduction drive also maternal effects on offspring fitness ex=. Our data revealed a paradoxical behaviour of *An*.*gambiae* whereby progeny of old females live longer, resist better to fungal infection but are more susceptible to *P. falciparum*. The fact that progeny of old females live longer is in sharp contrast to the dogma that offspring of old females display shorter life expectancy both in invertebrates and vertebrates (*30*) Further, our results with *P. falciparum* are also contrasting with the reported higher susceptibility to pathogen of progeny born from young females than those born from old females, in the *Daphnia* model system (*33*).

Epigenetics is a major area for contributing to understand maternal effects, aging processes as well as transgenerational acquired phenotypes (*35*). Both age and nutritional conditions of mothers have been implicated in epigenetic driven maternal effect in vertebrates (*34, 36*) and in invertebrates (*37–39*). Whereas epigenetic inheritance studies are expanding in *Drosophila* and *Caenorhabditis elegans*, little is known in *Anopheles* mosquitoes, notably at the molecular level (*4, 21, 25, 28, 40, 41*). Epigenetic investigation is therefore likely one leading area to develop in understanding the phenotypic results reported in this manuscript.

Regarding malaria epidemiology, our discovery has important implications for both malaria transmission modelling and vector control because it establishes an unexpected link between vector competence and the mosquito population age structure. Thus, older females, that are the most epidemiologically important (*11, 42*) are also producing offspring with enhanced vectorial capacity due to higher susceptibility and higher longevity. In nature, the proportions of *P. falciparum* infected females is usually not very high, therefore the discovered higher susceptibility of offspring from non-infected old females should guide further to strategy controlling these mosquito subgroups. This reinforced a strategy proposed to limit insecticide resistance among malaria vectors by specific targeting old females (*42*).This finding is also consistent with a potential adaptation trait of *P. falciparum* to *An. gambiae*. Indeed, infection of mosquito females with greater longevity (ie those born form old females) would provide the highest likelihood for transmission of the parasites to the human host. Our finding might also contribute explaining the resurgence of malaria transmission in seasonal transmission pattern. According to our finding, mosquitoes born from old uninfected females having survived the dry season will be highly susceptible to *P. falciparum* and thus might contribute efficiently to malaria transmission rebound.

## Acknowledgments

We wish to thank Richard Paul and Louis Lambrechts for their help with statistical analyses and fruitful comments on the manuscript, Nicolas Puchot for figure design, Agnès Zettor for her contribution to some *P. falciparum* mosquito infection, Jean Paul Latgé’s team for *Beauveria bassiana* experiments and the ICAReB platform for their constant on demand provision with human blood for *P. falciparum* cultures and mosquito infections.

## Contribution

- **Experimental Design:** CM, IT, CB
- **Perform the experiments:** CM, IT, MT, CT, ST, AL
- **Analyzed the data:** CM, IT, EB, CB
- **Wrote the manuscript:** CM, IT, CB

## Supplementary information

### Supplementary Information 1

#### Material and Methods

##### Mosquitoes

*Anopheles gambiae* Yaoundé strain (Tchuinkam et al., 1993) established at the Institut Pasteur in 1998 was produced under standardized laboratory conditions. Adults were reared at 26°C ±1, 80% RH with a 12:12h photoperiod. They had access *ad libidum* to 10% sucrose. Unless specified larvae were reared at 26°C ±1 with a 12:12h photoperiod and fed on Tretramin® baby fish powder till reaching the L2 stage and then on cat food. For colony maintenance, females were blood fed twice a week on an anesthetized rabbit, following French regulation for animals (Agreement #A 75-15-01). The production of the DP1 to DP4 offspring was as described in Figure 1. To accommodate different duration of larval development (Fig1A) the natural temperature gradient of the insectary was exploited with careful daily follow up. For experimental infection, females were separated from males 4 days post emergence and starved from sugar 12 hours prior infection with *P. falciparum*.

##### Production of P. falciparum gametocytes and mosquito infection procedure

Gametocytes from *P. falciparum* strain NF54 were produced as previously described (Mitri et al., 2009) using an automatic tipper table system (Ponnudurai et al., 1982). Fourteen days after gametocytogenesis induction, male gametocyte maturity was controlled by performing an exflagellation test; parasitemia and proportion of mature male and female gametocytes were determined on Giemsa stained thin smears. For mosquito infection, 10 ml culture were centrifuged at 2,000 rpm (805 g), and the pellet was suspended in an equal volume of AB human serum (EFS Rungis). The infected red blood cells (RBC) were placed on top of a 1:1 premix of fresh RBC (ICAreB, Institut Pasteur) and AB human serum, mixed gently and introduced inside a membrane glass feeder previously warmed at 37°C. All steps of this process were performed at 37°C. Mosquitoes were allowed to feed in the dark for 15 minutes. Fully engorged females were transferred into small cages and were provided with 10% sucrose. Mosquitoes were maintained for 7 days at 26°C, at which time their midgut was dissected in a 2% bromo-fluoresceine-PBS solution (SIGMA Aldrich). Stained oocysts were detected under a microscope at magnification 200X. A mosquito was considered infected if at least one oocyst was detected. Prevalence of infection was determined as the proportion of infected females among all dissected females. Intensity of infection corresponds to the mean oocyst load of infected females. Detailed information on each series of gametocyte culture is summarised in Sup Table 3.

##### An. gambiae infection with Beauveria bassiana

Assays were conducted with DP1 and DP4 female mosquitoes produced as described in Figure 1B. *Beauveria bassiana* strain I93-825 (a gift from J.-P. Latgé, Institut Pasteur) was grown on Sabouraud agar plates at 25°C during 12 days to obtain spores, as previously described (Boucias et al., 1988). Three to four days-old female mosquitoes were cold anesthetized and rolled on a spore-containing petri dish in such a way that mosquito whole body surface was in contact with the fungus spores. Contact last for 15 seconds. Pools of 50 mosquitoes of each DP1 and DP4 were then transferred into labelled carboard containers and maintained in an incubator at 26°, 80% HR with free access to 10% sucrose. Dead mosquitoes were count and removed every day. Two replicates of 50 mosquitoes per condition were performed. Analysis was performed using a Cox proportional hazards regression model fitted to the data using treatments (DP1 and DP4) as predictor terms (Cox, 1972).

##### Survival rates of uninfected An. gambiae DP1 and DP4 progenies

DP1 and DP4 female mosquitoes were produced according to Figure 1B. Four days after emergence, 2 pools of 50 females from each DP1 and DP4 progeny were placed in small cages at 26°C, 80%HR with free access to 10% sucrose. Dead mosquitoes were count and discarded every 3 or 4 days until all mosquitoes die. Two independent biological replicates were performed and analysed using a Cox proportional hazards regression model (Cox, 1972).

## Supplementary Information 2

**Sup Figure 1:**
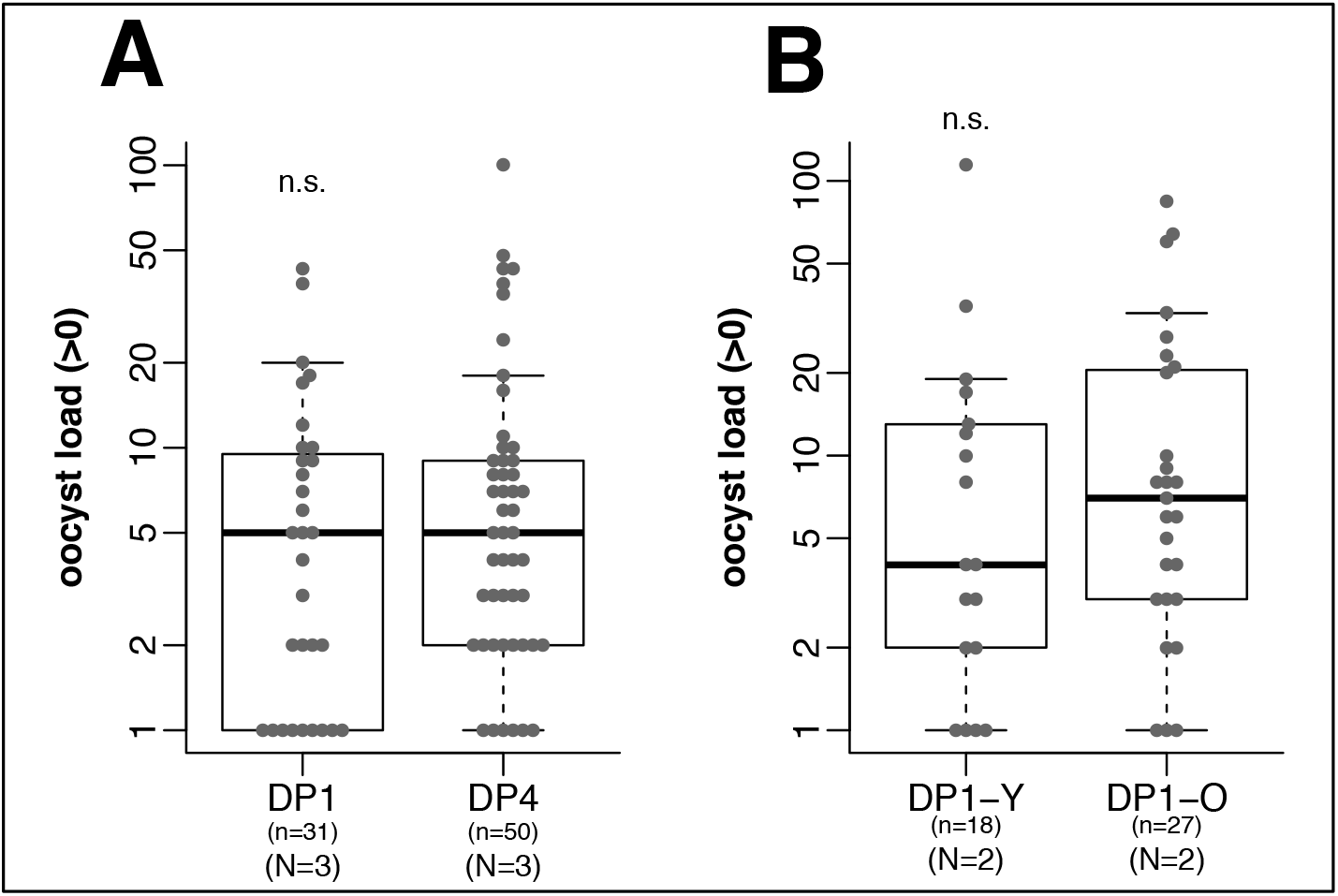
Oocyst development in offspring born from different gonotrophic cycles of mothers of the same age (Panel A) or from a first gonotrophic cycle from mothers of different age (Panel B) Panel A: DP1 and DP4 were produced as depicted in Figure 1 C and prevalence of infection are reported in Fig 4 A. **Panel B**: DP1 were produced form young or old females as depicted in Figure 1D and prevalence of infection are reported in Fig 4 B. Note than n in panel A and B takes into account only sample harboring at least on oocyst. Graph labels and statistical tests as in Fig 2 legend. Significance levels of adjusted Fisher-combined p-values: n.s., not significant; * p-value<0.05; ** p-value <0.01; *** p-value <0.001).

**Sup Figure 2:**
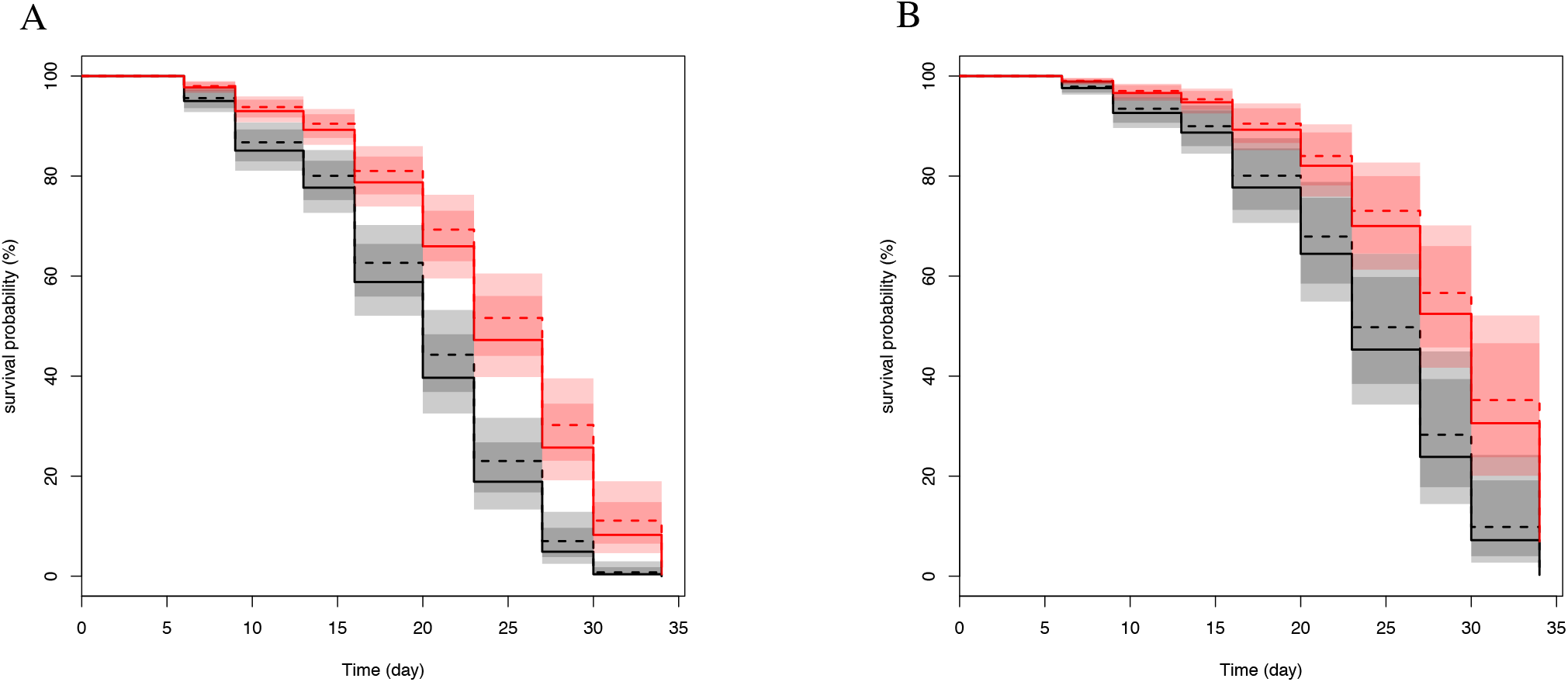
Differential survival rates of uninfected *An. gambiae* DP1 and DP4. Survival curves of *An. gambiae* DP1 (black) and DP4 (red). DP1 and DP4 were produced as depicted in Figure 1B. **A)** and **B)** are two biological replicates. For each biological replicate, two technical replicates were performed using 50 female mosquitoes in each replicate and condition (DP1/DP4). A Cox proportional hazards regression model was fitted to the data using condition and replicate as predictor terms. Taking into account all survival data, the p-value associated with condition (DP1/DP4) is 3.1e^−13^, and the p-value associated with the biological replicate is 3.0 e^−12^ while the p-value associated with the technical replicates is 0.21.

## Supplementary Information 3

**Supplemental Table 1:** P-values for pairwise comparisons for prevalence of P. falciparum infections between successive daughter populations (DP1 to DP4) born from the same mothers. Statistical differences in prevalence and intensity were first tested independently for each independent replication replicate using the Chi-Square test and p-values were empirically determined using 10^5 Monte-Carlo permutations. Following independent statistical tests, the p-values from independent tests of significance were combined using the meta-analytical approach of Fisher [1] when the direction of change of each independent replicate was concordant. Statistical tests have been done for all pairwise comparisons and p-values have been adjusted for multiple testing using the Bonferroni procedure. See also Figure 2A.

|  | DP1 | DP2 | DP3 |
| --- | --- | --- | --- |
| DP2 | 0.31 | NA | NA |
| DP3 | 0.034 | 1 | NA |
| DP4 | 1.8e-07 | 2.5e-05 | 0.003 |

**Supplemental Table 2:** P-values for pairwise comparisons for intensity of P. falciparum infections between successive daughter populations (DP1 to DP4) born from the same mother. Statistical differences in prevalence and intensity were first tested independently for each independent replication replicate using the Wilcoxon signed rank non-parametric test and p-values were empirically determined using 10^5 Monte-Carlo permutations. Following independent statistical tests, the p-values from independent tests of significance were combined using the meta-analytical approach of Fisher [1] when the direction of change of each independent replicate was concordant. Statistical tests have been done for all pairwise comparisons and p-values have been adjusted for multiple testing using the Bonferroni procedure. See also Figure 2B.

|  | DP1 | DP2 | DP3 |
| --- | --- | --- | --- |
| DP2 | 1 | NA | NA |
| DP3 | 0.81 | 0.59 | NA |
| DP4 | 0.0018 | 0.0016 | 0.0019 |

[1] Fisher RA. Statistical Methods for Research Workers. . Edinburgh: Oliver & Boyd; 1925. 356 p.

**Supplemental Table 3:** Parasitological characteristics of the *P. falciparum* gametocytes used for infecting *An. gambiae*.

| Data for |  | % gametocytes | % mature gametocytes | % male gametocytes | Exflagellation |
| --- | --- | --- | --- | --- | --- |
| Figure 2 | Exp1 | 2.4 | 71 | 46 | ++ |
|  | Exp2 | 1.7 | 66 | 60 | -/+ |
|  |  | 1.4 | 60 | 41 | ++ |
| Figure 3 | Exp1 | 2.4 | 83 | 60 | ++ |
|  | Exp2 | 1.3 | 100 | 28 | ++ |
|  |  | 1.5 | 100 | 50 | ++ |
|  | Exp3 | 1.8 | 79 | 57 | ++ |
|  |  | 1.7 | 83 | 64 | ++ |
|  |  | 1.6 | 84 | 54 | ++ |
|  | Exp4 | 1.7 | 70 | 55 | ++ |
|  |  | 1.5 | 86 | 56 | ++ |
|  | Exp5 | 1.6 | 92 | 61 | ++ |
| Figure 4A | Exp1 | 4.2 | 73 | 63 | ++ |
|  |  | 3.2 | 76 | 56 | ++ |
|  |  | 3.7 | 60 | 57 | ++ |
|  |  | 1.6 | 83 | 60 | ++ |
|  |  | 1.9 | 75 | 72 | ++ |
|  | Exp2 | 2.9 | 85 | 41.1 | +/- |
|  |  | 1.2 | 85.7 | 50 | +/- |
|  |  | 1.8 | 100 | 40.9 | ++ |
|  |  | 3.4 | 90.9 | 60 | +/- |
|  | Exp3 | 1.8 | 93.3 | 21.4 | ++ |
|  |  | 1.2 | 95 | 33.3 | ++ |
| Figure 4B | Exp1 | 3 | 93 | 50 | ++ |
|  | Exp2 | 0.72 | 100 | 55 | ++ |
|  |  | 1.62 | 91 | 54 | ++ |

The left column refers to the figure numbers for which culture were produced. Shaded data correspond to infection using a single culture flask. All the other infections were performed with pooled cultures for which the parasitological characteristics are provided

